# Exaptation of inactivated host enzymes for structural roles in orthopoxviruses and novel protein folds revealed by protein structure modeling

**DOI:** 10.1101/2022.11.22.517515

**Authors:** Pascal Mutz, Wolfgang Resch, Guilhem Faure, Tatiana G. Senkevich, Eugene V. Koonin, Bernard Moss

**Author notes:** These authors have contributed equally to this work (Pascal Mutz, Wolfgang Resch).

## Abstract

Viruses with large double-stranded DNA genomes appear to have captured the majority of their genes from the hosts at different stages of evolution. The origin of many virus genes is readily detected through highly significant sequence similarity with cellular homologs. This is the case, in particular, for virus enzymes, such as DNA and RNA polymerases or nucleotide kinases, that retain their catalytic activity after capture by an ancestral virus. However, a large fraction of virus genes have no readily detectable cellular homologs so that their origin remains enigmatic. We sought to explore potential origins of proteins of unknown provenance encoded in the genomes of orthopoxviruses, a thoroughly studied virus genus which includes major human pathogens. To this end, we used AlphaFold2, to predict the structures of all 214 proteins encoded by orthopoxviruses. Among the proteins of unknown provenance, structure prediction yielded a clear indication of origin for 14, along with validating several inferences previously made by sequence analysis. The major trend that emerges from these findings is the exaptation of enzymes from cellular organisms for non-enzymatic, structural roles in virus reproduction which is accompanied by disruption of catalytic sites and overall drastic divergence which precludes detection of homology at the sequence level. Among the 16 orthopoxvirus proteins found to be inactivated enzyme derivatives, are the poxvirus replication processivity factor A20, an inactivated derivative of bacterial NAD-dependent DNA ligase; major core protein A3, an inactivated deubiquitinase; F11, an inactivated prolyl hydroxylase; and more similar cases. However, for nearly one third of the orthopoxvirus virion proteins, no significantly similar structures were identified, suggesting exaptation with subsequent major structural rearrangement, yielding novel protein folds.

## Introduction

Viruses are ubiquitous, obligatory intracellular parasites of all life forms. Virus genome sizes vary over three orders of magnitude, from about two kilobases comprising a single gene to more than two megabases, with thousands of genes (1). Virus genes comprise three major functional classes: i) genes encoding components of virus replication machinery, ii) genes for structural components of virions and proteins involved in morphogenesis, iii) genes encoding proteins involved in virus-host interactions. The fractions of the genes in these classes depend on the size of the virus genome. Viruses with small genomes, in particular, most of the RNA viruses and all ssDNA viruses, primarily encompass genes of the first two classes, with few dedicated genes involved in interactions with the hosts. In contrast, in viruses with large dsDNA genomes, many of the genes are involved in various aspects of virus-host interactions, in particular, counter-defense. The first two gene classes include a small set of virus hallmark genes that are conserved in a broad range of viruses (2). Some of the virus hallmark genes encoding proteins involved in replication appear to originate in a primordial pool of genetic elements, whereas hallmark genes encoding major virion components can be traced to ancient acquisitions of cellular genes (3). For many other genes from all three functional classes, more recent cellular ancestry is readily traceable through significant sequence similarity with the apparent cellular ancestors. However, the provenance of numerous other virus genes remains obscure because no cellular homologs are detectable even with the most sensitive methods for protein sequence comparisons.

Protein structures are uniformly more strongly conserved in evolution than sequences so that structural comparison can illuminate the origin and function of many proteins that remain intractable at the sequence level. However, until very recently, the utility of structural comparison for the study of protein evolution remained severely hampered by the technical difficulty as well as time and labor cost of protein structure determination. The recent revolution in protein structure prediction ushered by the new artificial intelligence based methods, AlphaFold and RosettaFold, has dramatically expanded the opportunities for detecting homologous relationship among proteins by comparison of protein structure models to experimentally solved structure or other models (4, 5). For instance, recent benchmarking suggests that structural similarities now can be detected for half of the human proteins that have been considered “dark matter” (6, 7).

We were interested in exploring the potential of this new generation of protein structure prediction methods in uncovering the origins of the “dark matter” of virus genomes. We selected as the target a thoroughly studied group of viruses with large (about 200 kb) dsDNA genomes, the orthopoxviruses (ORPV), which include one of the most historically devastating human pathogens, variola virus, and the current major threat to human health, monkeypoxvirus, as well as vaccinia virus (VACV), one of the most popular model systems in virology (8, 9).

Altogether, ORPV possess 214 genes (OPG), of which subsets have been differentially lost in different virus lineages (10). Over the decades of study, each of these genes has been extensively analyzed, computationally, and for the most part, experimentally as well (8, 9). Nevertheless, for nearly half of the ORPV proteins, no homologs from cellular life forms could be detected even using the most sensitive methods of sequence analysis, and thus, the provenance of these genes remained enigmatic (10). We employed AlphaFold2 to predict the structures of all ORPV proteins and compared the resulting models to the databases of experimentally solved protein structures as well as pre-computed AlphaFold2 models. This analysis identified apparent cellular ancestors for 14 ORPV genes that until now belonged to the dark matter. An emerging major trend is exaptation of host enzymes for non-enzymatic, structural roles in virus reproduction, which is typically accompanied by disruption of the catalytic sites and extensive divergence such that homologous relationships become undetectable at the sequence level. However, the origins of many OPGs remain undecipherable even through structure comparison, suggesting emergence of multiple novel protein folds during poxvirus evolution.

## Results

### Structure predictions for orthopoxvirus proteins

Structure was predicted for representative sequences of each of the 214 OPG using AlphaFold2 (4). High quality models indicative of globular structure were obtained for 186 proteins (mean plddt score >= 70; Supplementary table S1). For additional 8 proteins, although the overall prediction was of comparatively low quality, individual globular domains had a mean plddt >= 70 (Supplementary table S1). These 194 high quality models of orthopoxvirus proteins were then compared to the PDB database of protein structures and to the AphaFold.db, the database of pre-computed models, using FoldSeek (very fast but relatively low sensitivity)(11) and Dali (relatively slow but higher sensitivity) (12). Similar structures with significant scores (e < e-03 for FoldSeek and/or z >5 for Dali; were detected for 188 proteins, and for 179 of these, the similar structures included cellular proteins, whereas for the remaining 9 only viral structures were retrieved(Supplementary table 1). All outputs were manually examined for the extent and quality of structure superposition and alignment, and for uncharacterized proteins without matches above the threshold, hits with lower scores were assessed. The known homologous relationships were accurately reflected by structure predictions. In addition, predictions with significant scores were also obtained for 14 OPGs, for which no homologs were previously detected, and these are considered in detail in the next section.

**Table 1.**
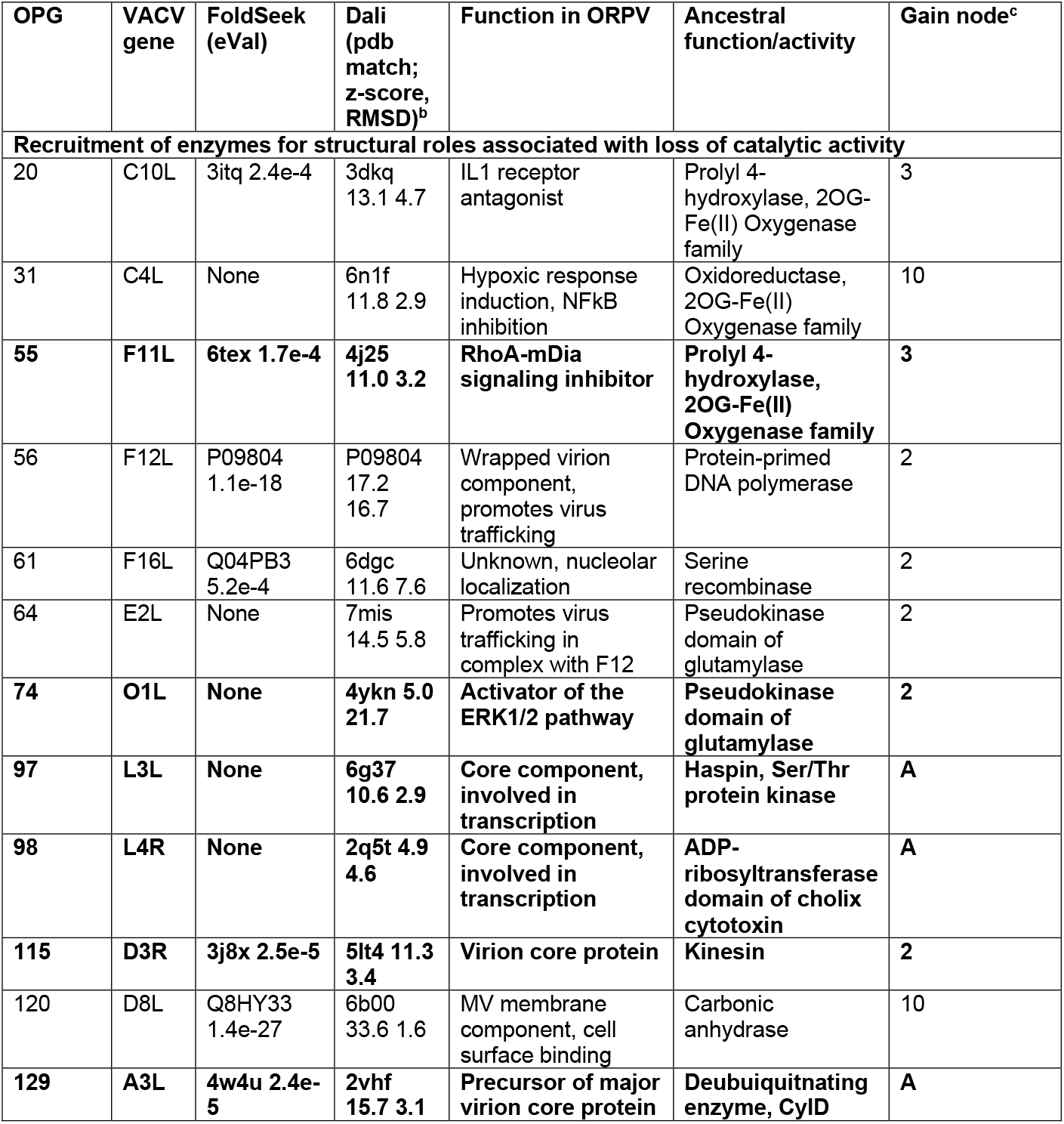
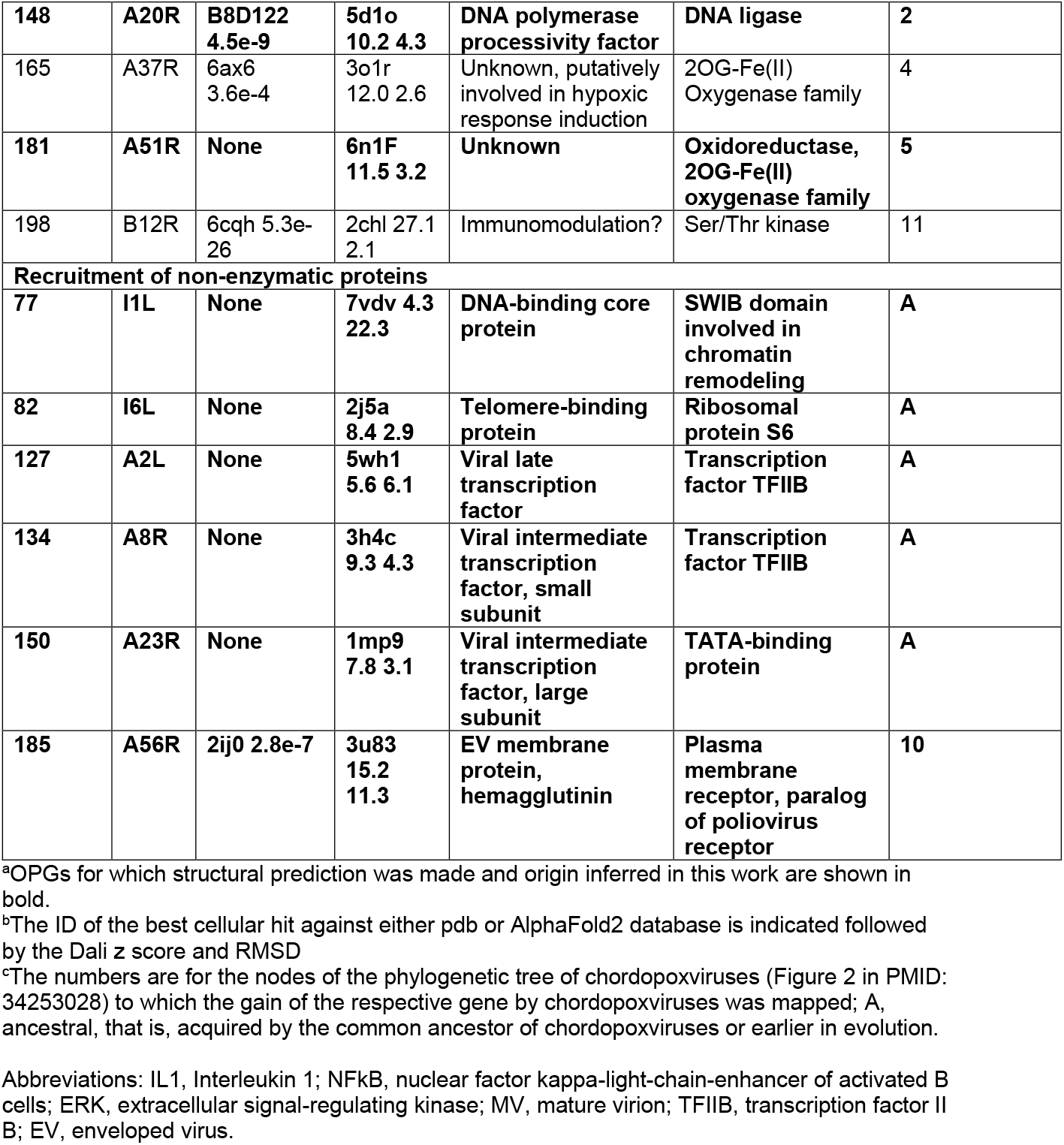
Exaptation of cellular proteins in orthopoxviruses^a^

### Exaptation of cellular proteins for viral functions

Our analysis of the structural models of ORPV proteins identified previously undetected likely origin from cellular ancestors for 14 OPG (Supplementary Table S1). Also, the results convincingly supported previous findings on exaptation of host enzymes for non-enzymatic functions in poxviruses that were originally made by using sensitive methods for sequence analysis (Table 1). In particular, OPG56 (F12L), which is involved in virus egress from infected cells (13), was shown to be a derived, inactivated DNA polymerase (14). Another poxvirus protein with yet unknown functions, OPG61 (F16L), is an inactivated serine recombinase (15). Three homologous OPGs were shown to be inactive prolyl hydroxylases, OPG20 (C10L), OPG31 (C4L) and OPG165 (A37R) (10). Structure modeling in this work fully validated the sequence-based inferences for these proteins, with high structure similarity scores (Supplementary table S1) and convincing structural superposition of the respective core domains (Figure 1, Figure S1).

**Figure 1.**
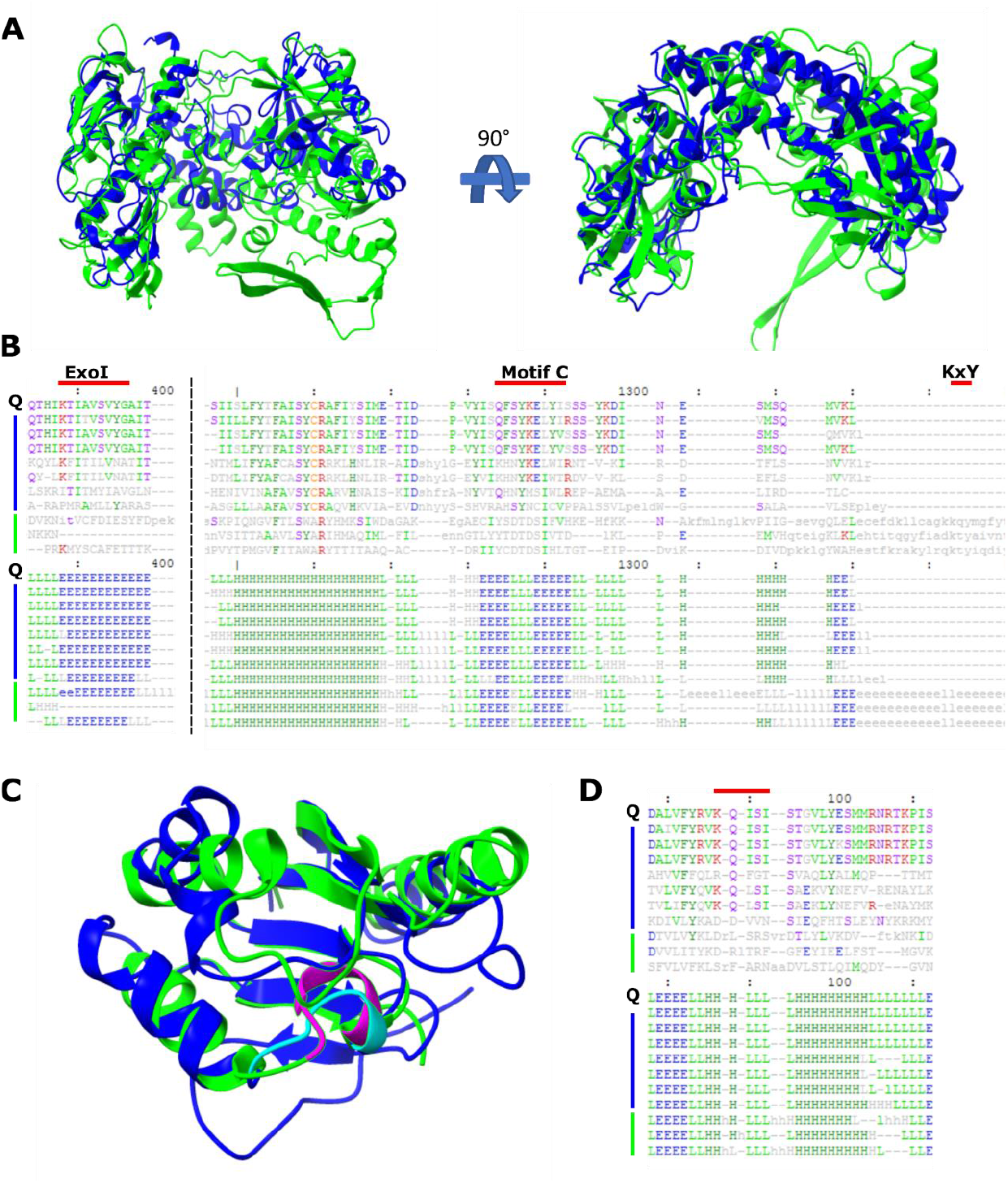
Structural modeling validates cases of enzyme exaptation discovered through sequence similarity. **A)** OPG56 (F12L) (blue, aa 205-634) and best Dali hit, a DNA polymerase type-B from yeast (AF-P09804-F, green, aa 338-916). **B)** Structural alignment of prototype OPG56 (Q), 7 OPG members from diverse chordopoxviruses (blue, from top to bottom: VARV, MPXV Zaire 96-I-16, VACV, SFV, MyxV, SORPV and MCV subtype 1) and three top hits found by Dali (green, DNA polymerases type-B from Kluyveromyces lactis (af2-db P09804), Claviceps purpurea (af2-db P22373) and Bacillus virus phi29 (2py5-B (60)). Alignment parts corresponding to ExoI motif, polymerase motif C and KxY motif are highlighted by red bar. Numbers indicate position in the structural alignment. **C)** OPG61 (F16L) (blue, aa 1-118 (of 231)) and best cellular Dali hit, the catalytic domain of a serin recombinase from Sulfolobus sp. L00 11 (archaea) (pdb 6dgc, (61) green, aa 65-164 (of 211)). Exemplified catalytic subdomain DRLXR (aa 139-143) in serin recombinase (magenta) and mutated stretch KQISI (aa 73-77) in OPG61 (cyan) are highlighted. **D)** Structural alignment of prototype OPG61 (Q), 7 OPG members from diverse chordopoxviruses (blue, from top to bottom:MPXV Zaire 96-I-16, VACV, VARV, MCV subtype 1, SFV, MyxV and SORPV)) and three Dali hits (green, an integrase from Lactococcus phage TP901-1 (3bvp-B (62)), an IS607-like serine recombinase from Sulfolobus sp. L00 11 (6dgc-D (61)) and a resolvase family site-specific recombinase from Streptococcus pneumoniae SP19-BS75 (3guv-A (63)). Red bar highlights catalytic center (DRLxR motif) of serin recombinases.

We identified 8 additional cases of apparent exaptation accompanied by the loss of enzymatic activity that were not detectable at the sequence level, bringing the total number of detected cases of exaptation of enzymes accompanied by inactivation in ORPV to 16 (Table 1 and Supplementary table S1). Two additional proteins, OPG55 (F11L) and OPG181 (A51R), were shown to be highly derived Fe-dependent dioxygenases, the protein superfamily that includes prolyl hydroxylases, three inactivated homologs of which were previously identified in chordopoxviruses as discussed above (Table 1 and Figure 2a,b).

**Figure 2.**
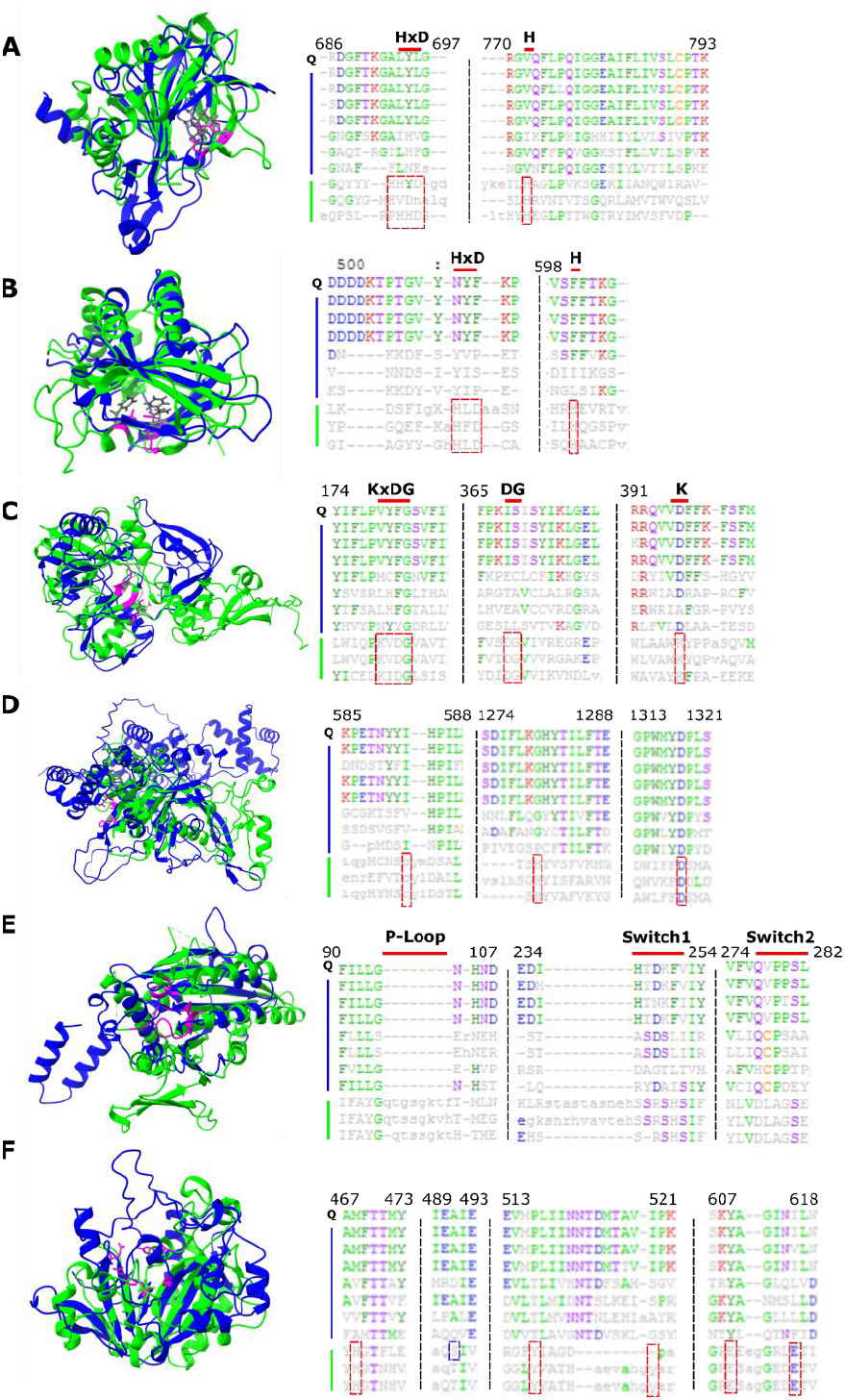
Newly identified cases of enzyme exaptation for structural roles in poxviruses accompanied by disruption of the catalytic sites. In each panel (A-F), the left subpanel shows superposition of the AlphaFold2 model of an OPG (blue) with a structurally similar cellular enzyme (green). Residues important for substrate binding and/or catalytic activity of the cellular enzyme are highlighted in magenta (cellular enzyme) and the corresponding residues within OPG are shown in grey. The right subpanel shows the structural alignment of OPG (Q), 7 OPG members from diverse chordopoxviruses (blue) and three structural homologs found by Dali (green). Proteins are listed from top to bottom. Catalytic and binding amino acid residues are highlighted in red. Numbers on top of the alignment refer to amino acid position in that alignment. **A) Left:** OPG55 (F11L) (aa 34-220) and human Lysyl Hydroxylase LH3 (6tex { DOI: 10.2210/pdb6tex/pdb }, aa 545 -738). Highlights: Residues known to bind Fe2+ and being essential for the catalytic activity within 2-OG dioxygenase enzyme members (H667, D669 and H719); OPG55 (L136, L138 and V184) are highlighted. **Right:** Structural alignment of OPG55 (Q), OPG55 from CMLV, VARV, MPXV Zaire-96-I-16, VACV, SwPV, SORPV ELK and LSDV NI-2490) and Dali hits (prolyl hydroxylase from Paramecium bursaria Chlorella virus 1 (5c5t-A (64)), PKHD-type hydroxylase from Psychrobacter sp. (af2-db A5WFM3) and human Lysyl Hydroxylase LH3 (6tex-A { DOI: 10.2210/pdb6tex/pdb })). **B) Left:** OPG181 (A51R) (aa 1-166) and Burkholderia pseudomallei oxidoreductase (6n1f { DOI: 10.2210/pdb6n1f/pdb }). Highlighted: Residues known to bind Fe2+ and being essential for the catalytic activity within 2-OG dioxygenase enzyme members (H134, D136, H188); OPG181 (N100, F102 and F150). **Right:** Structural alignment of prototype OPG188 (Q), OPG188 from VARV, CMLV, MPXV Zaire 96-I-16, YLDV, SwPV and LSDV NI-2490) and Dali hits (Oxidoreductase from Burkholderia pseudomallei (6n1f-B { DOI: 10.2210/pdb6n1f/pdb }), Fe2OG dioxygenase domain-containing protein from Dictyostelium discoideum (af2-db Q54K28) and Procollagen-proline 4-dioxygenase from Onchocerca volvulus (af2-db A0A2K6VMM0). **C) Left:** OPG148 (A20R) (aa 28-284) and a DNA ligase B from Klebsiella pneumoniae (af-db B5XTF0, aa 61-406). Highlights: Key amino acids of motifs I (KxDG), IV (DG) and V (K) within the ligase adenylation domain appear in the structure from left to right. **Right:** Structural alignment of prototype OPG148 (Q), OPG148 from VACV, MPXV Zaire-96-I-16, VARV, MyxV, Orf virus, MCV subtype 1 and CRV) and Dali hits (DNA ligases from Klebsiella pneumoniae (af2-db B5XTF0), E. coli (af2-db B7M4D2) and Streptococcus pneumoniae (af2-db B1IBQ3)). **D) Left:** OPG129 (A3L) and human CYLD USP domain (2vhf (65), aa 583 -955), a deubiquitinating enzyme. Highlights: Residues of the catalytic triad within the USP domain (C601, H871, D889); OPG129 (L136, L138 and V184). **Right:** Structural alignment of prototype OPG129 (Q), OPG129 from VACV, MyxV, VARV, MPXV Zaire-96-I-16, CRV, Orf virus and SGVP) and Dali hits (all CYLD USP domains found in: Danio rerio (af2-db, E7F1×5), Homo sapiens (2vhf-B (65)) and Sporothrix schenckii (af2-db, U7Q4Z6)). **E) Left:** OPG115 (D3R) and a kinesin motor ATPase from S. cerevisiae (1f9u (66), aa 385 – 722). Highlights: The P-loop (Walker A motif GxxxxGK(S/T)), Switch1 (SSRSH) and Switch2 (DLAGSE) motif within the ATPase. **Right:** Structural alignment of prototype OPG115 (Q), OPG115 from VACV, VARV, MPXV Zaire-96-I-16, MyxV, SFV, MCV subtype 1 and SOPV ELK) and Dali hits (all Kinesins from S. cerevisiae (1f9u-A (66)), Homo sapiens (5lt4-D (67)) and Drosophila melanogaster (5hnz-K (68)). **F) Left:** OPG98 (L4R) and best cellular Dali hit, Cholix toxin, a ADP-ribosyltransferase of *V. cholerae* (2q5t (69), aa 415 – 630). Highlights: The Cholix catalytic cluster (H460, Y493, Y504. E574, E581). **Right:** Structural alignment of prototype OPG98 (Q), OPG98 from VACV, VARV, MPXV Zaire-96-I-16, Orf virus, MyxV, MCV subtype 1 and CRV) and Dali hits (all ADP-ribosyltransferase toxins of: *P. aeruginosa* (af2-db: P11439) and *V. cholerae* (2q5t -A (69) and 3ki7-A { DOI: 10.2210/pdb3ki7/pdb) Residues of the catalytic cluster are highlighted in red. P11439 contains additional site highlighted in blue (S474).

Especially notable is the case of OPG148 (A20R), which showed highly significant structural similarity to DNA ligases, in particular, bacterial NAD-dependent ones (Table 1, Supplementary table S1 and Figure 2c). The VACV replisome consists of the DNA polymerase (OPG71, E9L) and two accessory subunits that function as processivity factors, OPG116 (D4R) and OPG148 (16, 17). Notably, both these proteins are derived from host enzymes involved in DNA replication and/or repair, namely uracil DNA glycosylase (UDG; OPG116) (18) and, as determined here, DNA ligase (OPG148). However, these proteins represent two contrasting modes of exaptation. OPG116 retains high similarity to eukaryotic UDGs and the corresponding enzymatic activity which is, however, not required for the function of this protein in VACV replication (19) although catalytic site mutations reduce replication in quiescent cells and attenuate virulence (20). Conversely, in OPG148, the ligase catalytic site is disrupted (Figure 2c), and the protein sequence diverged beyond recognition such that structure comparison was essential for inferring the origin of this virus protein.

Another ORPV protein, OPG129 (A3L), a major core protein (21), is an inactivated derivative of a distinct deubiquitinating enzyme (DUB) (Figure 2d). Recruitment of DUBs as proteases catalyzing processing of virus proteins is a well-known phenomenon exemplified by OPG83 (I7L), the protease involved in virion protein processing in poxviruses and many other viruses of the realm *Varidnaviria* (22). We are unaware of previously reported cases of exaptation of DUBs for structural roles although in alphaviruses, a serine protease has been exapted as the major capsid protein (23).

Another newly identified case of enzyme exaptation for a structural role in orthopoxviruses involves recruitment of a major cytoskeleton component. OPG115 (D3R), a virion core protein (24), appears to be a derivative of the enzymatic domain of kinesin, the motor ATPase that is involved in various intracellular trafficking processes (Figure 2e). Recapitulating the pattern observed in other enzymes exapted for structural roles, the catalytic residues of the ATPases, in particular, the canonical Walker A and B motifs are replaced in OPG115 (Figure 2c).

OPG98 (L4R), a core protein involved in VACV early transcription, showed structural similarity to the C-terminal ADP-ribosyltransferase domain of *Vibrio cholerae* Cholix cytotoxin that additionally contains an N-terminal receptor recognition domain (25). Because of the presence of the N-terminal domain, the Dali score in this case was relatively low, but OPG98 fully superimposed over the ADP-ribosyltransferase domain whereas the catalytic residues are replaced (Figure 2f).

Four OPGs contain inactivated protein kinase or pseudokinase domains. OPG97 (L3L) is a VACV core component involved in the transcription of early genes. The AlphaFold model of this protein showed significant structural similarity to Ser/Thr proteins kinases, in particular the atypical kinase domain of haspin, an animal chromatin remodeling regulator (26), but the catalytic site residues are partially replaced in the poxvirus protein (Figure 3a). Notably, the aspartate in the active site has not been exchanged for crododilepox virus OPG97 but the lysine at the ATP-binding site is present. Conversely, in the OPG97 proteins from other chordopoxviruses, the lysine at the binding site is intact whereas the aspartate was mutated. Notably, OPG97 is a second inactivated protein kinase in chordopoxviruses, along with OPG198 (B12R) that has a much higher similarity to active Ser/Thr kinases, readily detectable at the sequence level, again, with only partial replacement of the catalytic residues (Figure 3a), which apparently reflects a later acquisition of a host kinase (see below). The two inactivated kinase derivatives followed different paths of exaptation, namely, recruitment for an essential structural role in the case of OPG97 and apparent involvement in immunomodulation in the case of OPG198 (27, 28).

**Figure 3.**
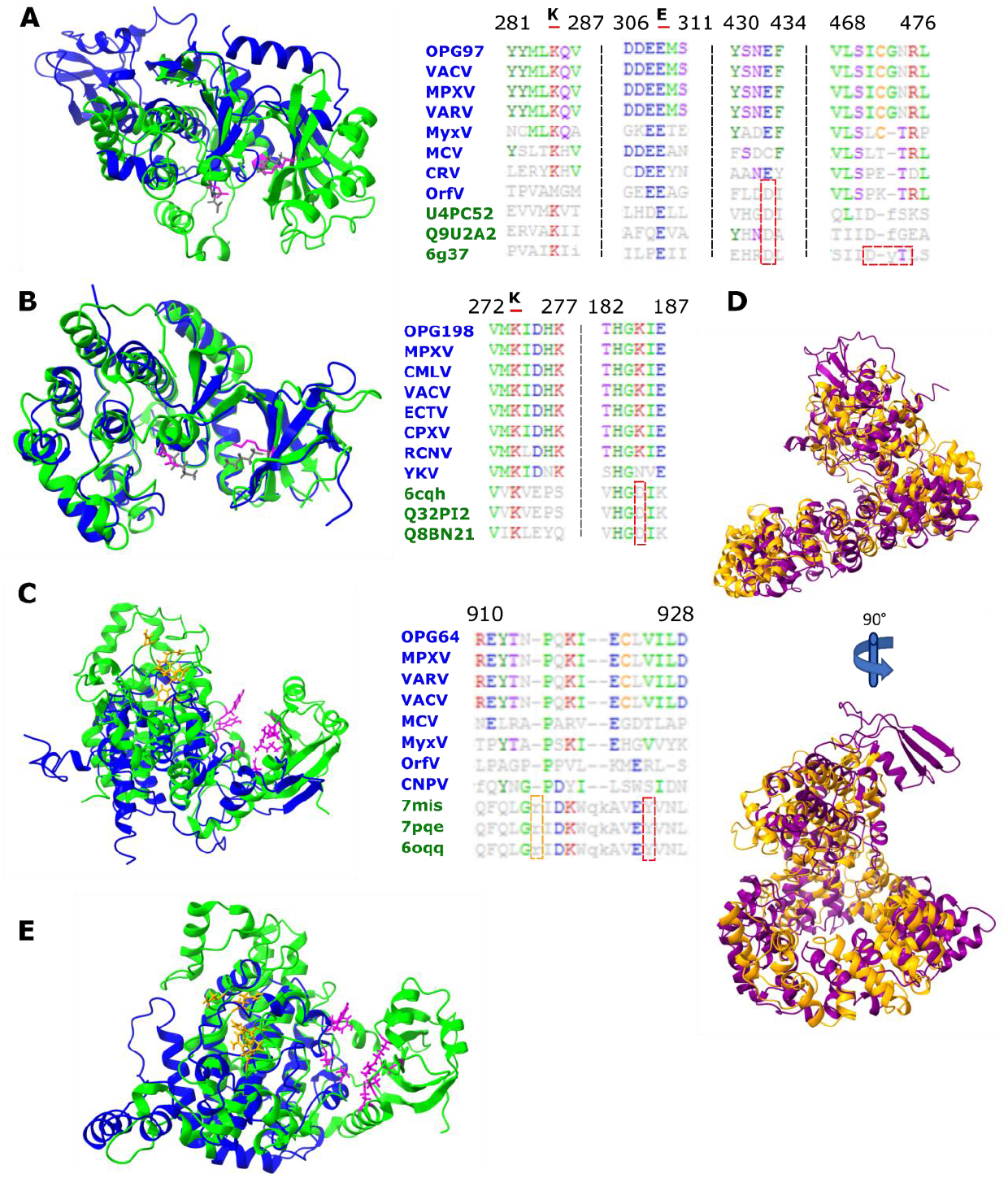
Inactivated kinases and pseudokinases in orthopoxviruses. **A) Left:** OPG97 (L3L) (blue, aa 66 – 350) and Haspin, an atypical Ser/Thr kinase (green, 6g37 {Heroven, 2018 #2959},aa 472-798); (Mutated) ATP binding site, helix αC glutamate and active site are highlighted (K511, E535 and D649 in Haspin, magenta; K93, E99 and E177 in OPG97, grey). **Right:** Structural alignment of prototype OPG97, 7 OPG members from diverse chordopoxviruses and three Dali hits, all kinases. Haspin specific ATP-binding motif DYT is highlighted in red. PDB structure: 6g37 (70). **B) Left:** OPG198 (B12R) (blue) and human vaccinia-related kinase (VRK, 6cqh {DOI: 10.2210/pdb6cqh/pdb}, green, aa 22-341). ATP binding site and active site are highlighted (K71 and D171 in VRK, magenta and K45 and K139 in OPG198). **Right:** Structural alignment prototype OPG198 (Q), 7 OPG members from diverse chordopoxviruses and three Dali hits, all vaccina-related kinases. PDB structure: 6cqh {DOI: 10.2210/pdb6cqh/pdb }. **C) Left:** OPG64 (E2L) (blue, aa 444-737) and best cellular hit, SidJ, a glutamylation protein with pseudokinase-fold from Legionella pneumophila (7mis (30), green, aa 336-758). Key amino acids of SidJ nucleotide-binding pocket (H492, R500, Y506, R522, N733 (orange)) and SidJ kinase-like active site (R352, K367, E373, E381, Y452, Y532, N534, D542 (magenta)) are shown. **Right:** Structural alignment of prototype OPG64 (Q), 7 OPG members from diverse chordopoxviruses (blue) and three Dali hits (green). Residues important within SidJ for nucleotide binding (R522, orange) and kinase-like activity (Y532, red) are highlighted. PDB structures: 7mis (30), 7pqe (29), and 6oqq (71). **D)** Superposition of OPG64 (purple) and OPG74 (O1L) (orange). **E)** Pseudokinase domains of OPG74 (blue, aa 380 - 666) and SidJ (7mis (30), green, aa 336 -758). Sites highlighted as in C).

Two more inactivated kinase homologs contain domains structurally similar to the pseudokinase domain of bacterial glutamylases SidJ in which the pseudokinase domain catalyzes ATP-dependent glutamylation that, in the case of *Legionella pneumophyla*, inactivates bacterial ubiquitin ligase (29, 30). OPG64 (E2L) is a large, two-domain protein for which the structure was recently solved and shown to consist of a globular head domain and an annular (ring) domain comprised of multiple α-helices (30, 31). The head domain was found to share structural similarity with the pseudokinase domain of SidJ but this similarity was interpreted as weak and potentially spurious (31). In our comparison, OPG64 produced a highly significant Dali score with SidJ (Table 1), and the overlay of the two domains included superposition of all core elements of the pseudokinase domain (Figure 3b), strongly suggesting that the head domain of OPG64 is indeed derived from the pseudokinase. However, the catalytic residues of the pseudokinase domain are replaced in OPG64 (Figure 3b), following the trend of inactivation of exapted enzymes. The annular domain of OPG64 seems to be unrelated to any structure outside of the ORPV. We further observed that another large ORPV protein protein, OPG74 (O1L), showed significant structural similarity to OPG64, with full superposition of both domains (Figure 3c) although the similarity of the OPG74 head domain to the pseudokinase was much lower than in the case of OPG64 (Figure 3b,d). The conservation of the domain architecture, including the unique annular domain, implies that, notwithstanding the low sequence similarity, OPG64 and OPG74 evolved as a result of ancient gene duplication in an ancestral chordopoxvirus, followed by extensive divergence. Despite the structural similarity and apparent common origin, OPG64 and OPG74 perform quite different roles in ORPV: OPG64 is involved in trafficking in infected cells (32) whereas OPG74 activates the ERK1/2 pathway and promotes virulence (33). Notably, OPG64 functions as a complex with OPG56 (F12L) (32), another inactivated enzyme, a derived DNAP discussed above.

Six other cases of apparent exaptation of cellular proteins for viral functions involve non-enzymatic, structural proteins (Table 1; Supplementary table S1; Figure 4). The poxvirus telomere-binding protein, OPG82 (I6L) (34), appears to be a derivative of the ribosomal protein S6. In this case, recruitment of a nucleic acid-binding protein might not involve a radical functional change. Even less surprising seems to be the adoption of the transcription factor TFIIB as poxvirus late (OPG127 (A2L)) and intermediate (OPG134 (A8R)) transcription factors (35, 36). Notably, OPG77 (I1L), a DNA-binding core component, was shown to contain a SWIB domain that is present in various chromatin proteins and is involved in chromatin remodeling (37); a simlar role of this domain in the poxvirus core can be envisaged. Along the same lines, OPG150 (A23R), an intermediate transcription factor (38), shows similarities to a TATA binding protein (1mp9), using the primary function of the original protein, DNA binding, during poxvirus life cycle. OPG185 (A56R), an ORPV membrane protein and hemagglutinin (39), appears to be a derivative of a cellular receptor.

**Figure 4.**
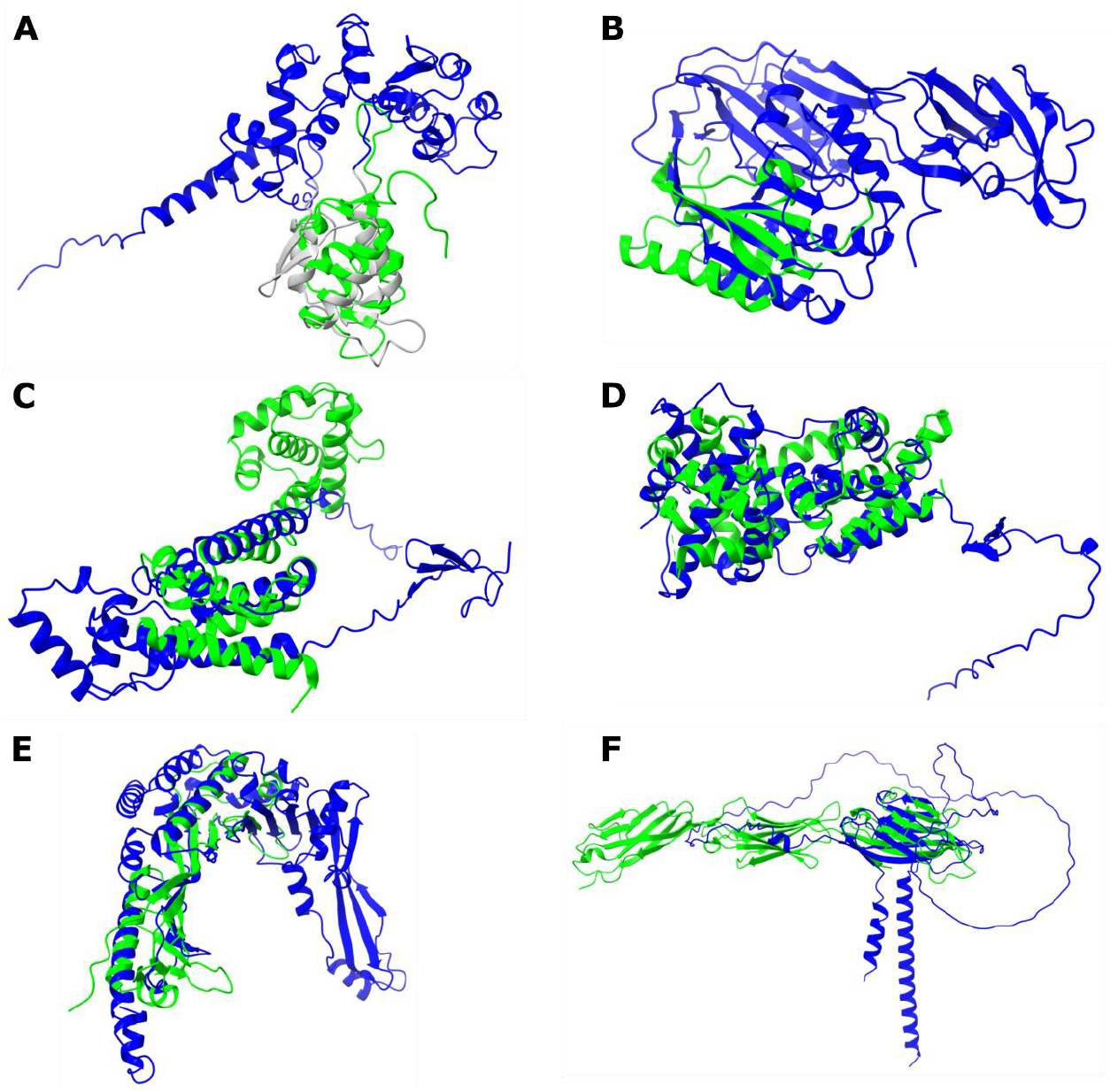
Newly identified cases of exaptation of non-enzymatic proteins for structural roles in poxviruses. Superimposition of OPG models (blue) over the best structural match (green) icdentified by Dali. **A)** OPG77 (I1L) and SWIB domain of mouse BRG1-associated factor 60a in Mus musculus (1uhr { DOI: 10.2210/pdb1uhr/pdb }). The putative SWIB domain in OPG77 (amino acid positions 138-222) is rendered in grey. Mouse SWIB domain contains 2 small antiparallel beta-sheets. **B)** OPG82 (I6L) and ribosomal protein S6 (2j5a (72), Aquifex aeolicus). **C)** OPG 127 (A2L) and C-terminal region of Transcription Factor IIB (5wh1 (73), Homo sapiens). **D)** OPG134 (A8R) and C-terminal Domain of Transcription Factor IIB (3h4c (74)). **E)** OPG150 (A23R) and TATA-binding protein (1mp9 (75), Sulfolobus acidocaldarius). **F)** OPG185 (A56R) and nectin-1 (3u83 (76), Homo sapiens).

From the previous reconstruction of chordopoxvirus evolution, information on the stage of evolution at which each of the exapted genes was captured by the viruses can be extracted (10). The majority of these genes were gained at an early stage of chordpoxvirus evolution along the branch separating fish poxviruses from the rest of the chordopoxviruses (Table 1). The exceptions are OPG31 (C4L) and OPG198 (B12R) that emerged at a later stage of chordopoxvirus evolution, the first one via duplication of OPG20 (an already inactivated prolyl hydroxylase) and the second one by capture of an ancestral protein kinase from the host. The multiple alignments of all exapted enzymes show consistent replacement of the catalytic amino acid residues in all poxviruses (Figure 2) demonstrating that the respectrive enzymatic activities were lost shortly after the capture of the respective host genes by the ancestral poxviruses. An apparent exception to this pattern of enzyme inactivation is OPG61 (F16L), where the deepest poxvirus branch carrying this gene, the crocodilepoxviruses, appears to encode an active serine recombinase, with inactivation apparently having occurred along the branch separating crocodilepoxviruses from the rest of chordopoxviruses (15).

### Routes of evolution of orthopoxvirus genes

Recently, the virus genes acquired from the hosts have been classified into five categories with respect to the degree of functional change of the encoded proteins (40):

1. Virus Hallmark Proteins that were acquired at the earliest stages of virus evolution (like capsid proteins) or possibly inherited from primordial replicators (like some replicative enzymes)
2. “Radical” exaptation accompanied by major change in the protein function
3. “Conservative” exaptation of host proteins when the original activity is exploited for virus functions
4. Direct exploitation of host proteins in their original capacity
5. Virus proteins of unknown provenance, some possibly evolving de novo.

Furthermore, a distinction has been made between “extramural” exaptation that involves direct recruitment of host genes and “intramural” exaptation when proteins encoded by the virus itself are repurposed either via a duplication or functional moonlighting (40).

The structural comparisons presented allow us to reach greater confidence in inferring the likely origins of viral proteins than previously attainable. Here we classified the OPGs according to the above categories (Table 2; Figure 5). There are only three bona fide hallmark genes in poxviruses, those encoding the homolog of the major capsid protein involved in virion morphogenesis, the packaging ATPase and a single replicative enzyme, the primase-helicase.

**Table 2.** Inferred routes of evolution of orthopoxvirus proteins

| Evolutionary/functional category | Number of genes | OPG (genes)<sup>a</sup> |
| --- | --- | --- |
| Virus Hallmark Genes | 3 | 125(D13L), 160 (A32L), 117 (D5R) |
| Radical exaptation, enzymes recruited for structural roles | 18 | 20 (C10L), 31 (C4L), **55 (F11L)**, 56 (F12L), 61 (F16L), **64 (E2L)**, **74 (O1L)**, **97 (L3R)**, **98 (L4R)**, **115 (D3R)**, 116 (D4R), 120 (D8L), 129 (A3L), 148 (A20R), 165 (A37R), 175 (A45R)<sup>b</sup>, 181 (A51R), 198 (B12R) |
| Conservative exaptation, proteins recruited for new function based on the same activity | 95 | 2-6 (C23L-16L), 8-11, 13-17, 19 (C11R), 21-23, 25 (C9L), 29 (C6L), 30 (C5L), 32-34 (C3L-1L), 35-36 (N1L-2L), 37 (M1L), 39-41 (K1L-3L), 44 (K7R), 45 (F1L), 47 (F3L), 49 (F5L), 54 (F10L), 57 (F13L), 63 (E1L), 65 (E3L), 67 (E5R), 72 (E10R), 83 (I7L), **82 (I6L)**, 84 (I8R), 85 (G1L), 89 (G5R), 91 (G6R), 93 (G8R), 106 (H1L), 108 (H3L), 113 (D1R), 118 (D6R), 121-124 (D9R, D10R, D11L, D12L), 133 (A7L), 145 (A18R), 146 (A19R), **150 (A23R)**, 159 (A31R), 161 (A33R), 162 (A34R), 167-169 (A38L-40L), 176 (A46R), 177 (A47L), 179 (A49R), 182-**185 (A52R-56R)**, 187-191 (B1R-6R), 193 (B8R), 194<sup>c</sup>, 196 (B10R), 199-201 (B13R-16R), 203-205 (B18R-20R), 206, 208 (C12L), 210<sup>c</sup>, 211 (C15L), 212-214 |
| Direct functional recruitment | 30 | 42 (K4L), 43 (K5L, K6L), 46 (F2L), 48 (F4L), 66 (E4L), 71 (E9L), 75 (O2L), **77 (I1L)**, 79 (I3L), 80 (I4L), 88 (G4L), 90 (G5.5R), 101 (J2R), 102 (J3R), 103 (J4R), 105 (J6R), 109 (H4L), 111 (H6R), 119 (D7R), **127 (A2L)**, 131 (A5R), **134 (A8R)**, 149 (A22R), 151 (A24R), 156 (A29L), 171 (A42R), 174 (A44L), 178 (A48R), 180 (A50R), 186 (A57R) |
| Origin unknown | 68 | 1, 7, 12, 18, 24, 26 (C8L), 27 (C7L), 28, 38 (M2L), 50-53 (F6L-9L), 58 (A14L), 59 (A14.5L), 60 (F15L), 62 (F17R), 68-70 (E6R-8R), 73 (E11L), 76 (O3L), 78 (O2L), 81 (I5L), 86 (G3L), 87 (G2R), 92 (G7L) 94 (G9R), 95-96 (L1R-2R), 99 (5R), 100 (J1R), 104 (J5L), 107 (H2R), 110 (H5R), 112 (H7R), 114 (D2L), 126 (A1L), 128 (A2.5L), 130 (A4L), 132 (A6L), 135-139 (A9L-13L), 140 (A14L), 141 (A14.5L), 142 (A15L), 143 (A16L), 144 (A17L), 147 (A21L), 152-155 (A25L-28L), 157 (A30L), 158 (A30.5L), 163 (A35R), 164 (A36R), 166, 170 (A41L), 172 (A43R), 173 (A43.5R), 192 (B7R), 195 (B9R), 197 (B11R), 202 (B17L), 207 (C11.5R), 209 (C13L, C14L), |

<sup>a</sup>OPG numbers are indicated, with VACV-Copenhagen gene names given in brackets if available. The OPGs for which structure was predicted and origin inferred in this work are in bold.
<sup>b</sup>OPG116 (uracil DNA glycosylase) and OPG175 (superoxide dismutase) are special cases of exaptation where the enzymatic activity is retained but the principal role of the protein in ORPV is structural.
<sup>c</sup>OPGs194 and 210 are multidomain membrane spanning proteins with one domain resembling Ig protein superfamily. Most of the protein is modeled with low confidence and gives no convincing match.

**Figure 5.**
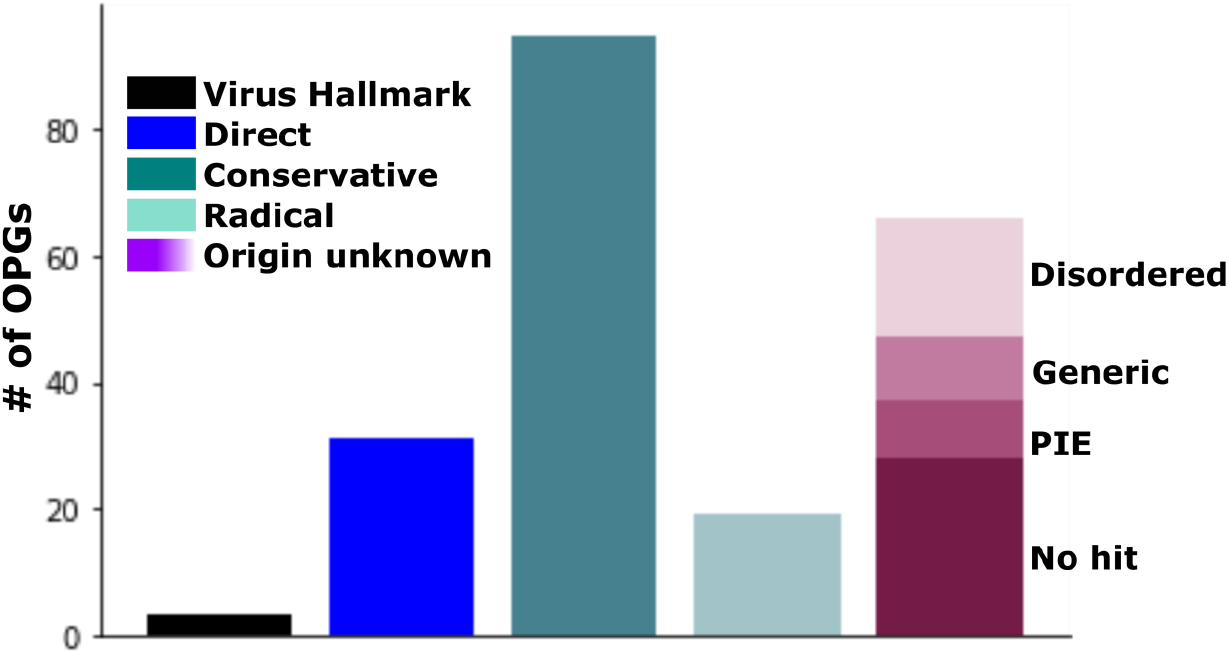
Inferred routes of evolution of orthopoxvirus proteins. The number of OPGs assigned to different classes of virus proteins with respect to the degree of functional change from the respective cellular ancestors are shown. Black, Virus Hallmark Proteins; blue, direct functional recruitment; blue-grey, “conservative” exaptation; opal, “radical” exaptation; shades of purple, unknown provenance. OPGs of unknown provenance were classified into disordered; those that were predicted to adopt a globular fold, but had no convincing match, with only generic matches (for example, to various β–sandwiches); PIE domains; predicted globular proteins with no match (mostly short proteins) (see Supplementary table S1 for details).

The inactivated enzymes adopted for structural roles by the viruses, as discussed in the preceding section, represent clear cases of radical exaptation. This route of exaptation involves major sequence and even structural changes to the proteins involved, which make the recognition of the ancestral relationships a non-trivial task. A distinct exaptation scenario is apparent for UDG, OPG116 (D4L), and superoxide dismutase,OPG175 (A45R) (41), which retain the catalytic sites and activity as well as high sequence similarity to their respective cellular homologs, and perform dual roles in ORPV reproduction, both structural and enzymatic. Conversely, one of the inactivated protein kinases discussed in the previous section, OPG198 (B12R), and OPG120 (D8L), an inactivated carbonic anhydrase, are highly similar to the respective cellular homologs despite the loss of the enzymatic activity, having been acquired relatively late in chordopoxvirus evolution (42).

The largest group of poxvirus proteins appears to have evolved via “conservative” exaptation (Table 2). This group includes most of the proteins involved in virus-host interactions, in particular, the three major families of paralogous poxvirus proteins, those containing ankyrin and PRANC domains, Kelch and BTB domains, and Bcl2 domains (10, 43). The functional diversification of virus proteins within these families represents “intramural exaptation” whereby virus proteins adopt a new function after gene duplication within evolving virus genomes.

The smaller group of poxvirus proteins that represent direct recruitment of cellular activities primarily consists of enzymes involved in genome replication and expression as well as those of nucleotide metabolism (Table 2).

Finally, one third (66) of the poxvirus proteins, particularly, virion structural components and proteins involved in virion morphogenesis, showed no convincing structural similarity to any available structures or AlphaFold2 models, and their origin thus remains obscure. Several of these are small, apparently, non-globular proteins, for which no good models were obtained, for example, OPG24 or OPG173 (A43.5R), and several others are tiny membrane proteins (OPG28, 59 (F14.5L), 76 (O3L), 78 (I2L)). However, more than half of the proteins in this group are globular, and the quality of their AlphaFold2 models is on average close to that for proteins with recognizable folds (mean plddt values of 83.3 and 86.1, respectively; Supplementary table S1, Figure S2). Some of these proteins did have a Dali match(es) with *z* score >5. However, the “grey zone” of Dali searches is wide (12), and inspection of matches for these proteins showed lack of consistency among the matching structures and/or large RMSD (Supplementary table S1), indicating lack of evidence of homologous relationships. Given that all models in this work were compared both to the PDB and to the large database of AlphaFold2 models that covered the complete proteomes of humans and other model organisms (44), the lack of similar structures strongly suggests that these proteins adopt unique folds that are missing or are extremely rare in cellular organisms. Of note, 10 proteins in this group are subunits of the entry-fusion complex (EFC) (45, 46); OPGs 53 (F9L), 86 (G3L), 94 (G9R), 95 (L1R), 99 (L5R), 104 (J5L), 107 (H2R), 143 (A16L), 147 (A21L), 155 (A28L), Figures S3 and S4) with OPG143 (A16L), OPG94 (G9R) and OPG104 (J5L) as well as OPG53 (F9L)-OPG95 (L1R) being known paralogs (46). Two additional pairs of paralogous proteins with apparent novel folds were detected: OPG18 (missing in VACV)-OPG27 (C7L) and OPG152 (A25L)-OPG153 (A26L). All these paralogous relationships among OPGs were validated by an all-against-all comparison of the AlphaFold2 models (Figure S4), but no additional significant structural similarities were identified, emphasizing the diversity of the OPG of unknown provenance, with apparent novel folds. Notably, in addition to the structural proteins, this group includes the fourth family of chordopoxvirus paralogous proteins, those containing the chemokine-binding PIE domain, an all-beta domain with a unique topology (47). Figure 6 illustrates 8 apparent unique folds, each representing a compact, globular structure with a high confidence prediction, at least, for the corresponding core domains.

**Figure 6.**
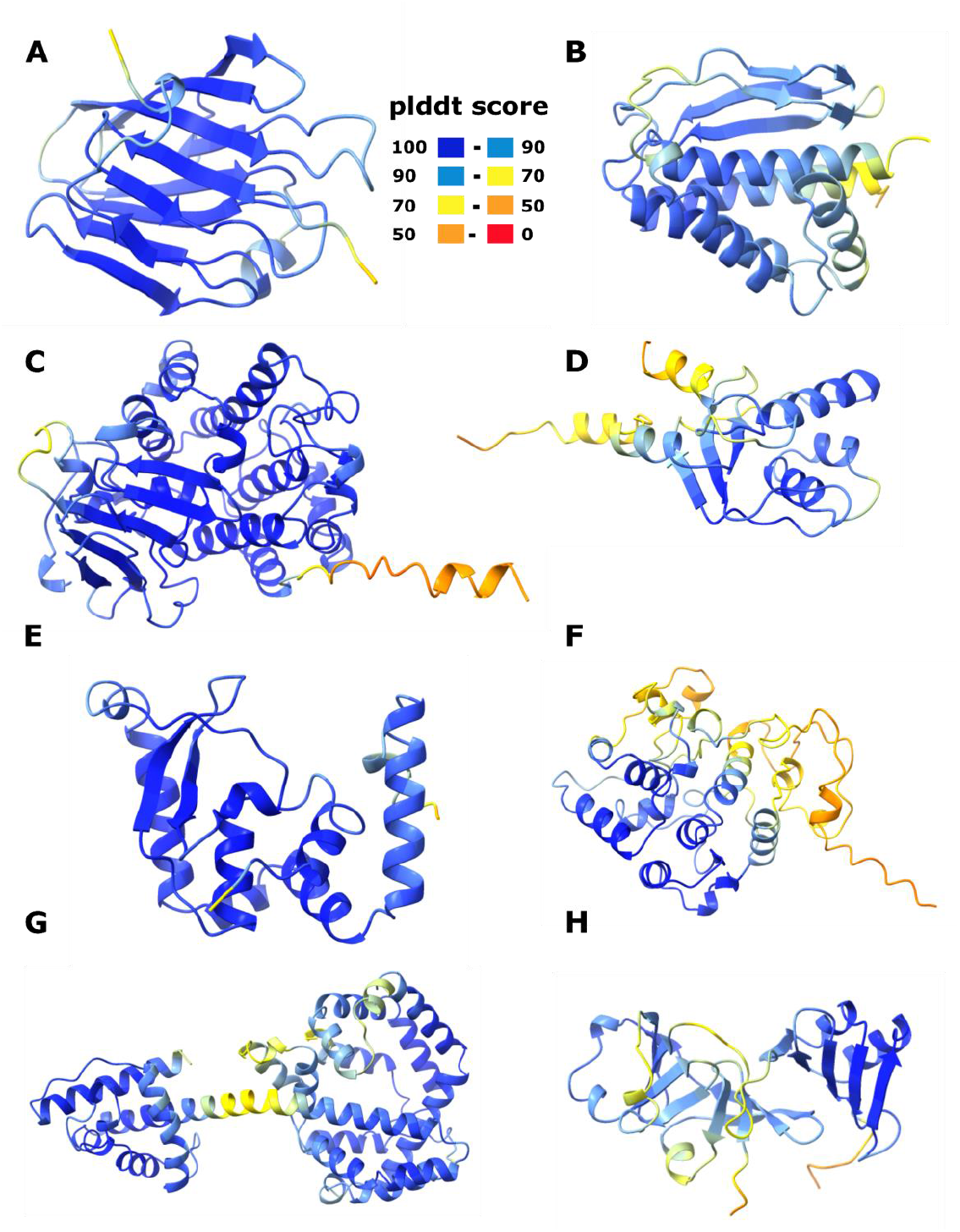
Predicted novel folds of orthopoxvirus proteins. Predicted globular structures of ORPV proteins with no homologs detrected outside of poxviruses are shown. Coloring according to AlphaFold2 plddt-score as shown in A). Weakly supported C-terminal domains not shown for OPG95 (L1R) and OPG153 (A26L). **A)** OPG27 (C7L), **B)** OPG95 (L1R) (aa 1-176 (of 250)), **C)** OPG153 (A26L) (aa 1-359 (of 518)), **D)** OPG114 (D2L), **E)** OPG112 (H7R), **F)** OPG70 (E8R), **G)** OPG132 (A6L) and **H)** OPG163 (A35R). OPG27, 95 and 153 have homologs among other poxvirus OPGs (Figure S4).

## Discussion

The recently developed methods for protein structure modeling, in particular, AlphaFold2 (4, 48), as well as RosettaFold (5), open up unprecedented opportunities for tracing origin of proteins through structural similarity. These developments are particularly promising for the study of the origins of virus proteins because viruses typically evolve (much) faster than cellular organisms (49, 50). We applied AlphaFold2 structural modeling to the proteins of ORPV, a well characterized group of large mammalian viruses of major medical importance (8, 9). High quality structural models were obtained for the great majority of the ORPV proteins, and for 14 proteins without any homologs detectable by sequence similarity, structural similarity pointed to the likely cellular ancestors. These findings include both radical exaptation, where a host enzyme is repurposed for a structural role in virus reproduction, whereas the catalytic activity is lost, and conservative exaptation where the repurposing involves the original activity of a protein (40). Exaptation of enzymes accompanied by disruption of the catalytic sites was detected for 8 additional ORPV proteins, bringing the total number of such cases to 16 (Table 1). These inactivated enzyme derivatives perform various functions in ORPV reproduction, but particularly notable is exaptation of enzymes for the role of major virus core proteins of which 4 cases were detected. It should be noted that, even the inactivated enzyme derivatives among the ORPV proteins display a broad range of similarity to the inferred cellular ancestors, from high sequence conservation (despite catalytic site disruption) to moderate (even if significant) structural similarity. Apparently, this broad spectrum of conservation reflects different stages on the evolutionary path of exaptation and different degrees of functional change. Taken together, the results emphasize the importance of both radical and conservative modes of exaptation in virus evolution and the utility of structural comparison in detecting this phenomenon.

However, apart from the identified cases of exaptation of cellular proteins, the surprising outcome of this work is that, for nearly one third of the OPGs, no similar structures that appeared indicative of the likely origin were detected. AlphaFold2 modeling (along with RosettaFold) followed by structure comparison is by far the most powerful current approach for detecting homologous relationships among proteins (51, 52). Further methodological improvement, certainly, will follow. However, given the high model quality already attained, together with the completeness of the databases used for structure comparison, with the PDB and AlphaFold2 predictions for multiple model proteomes combined, a radical improvement in the recognition of similar structures hardly can be expected. Hence the question of the origin of a large fraction of virus proteins becomes pressing. Several proteins of unknown provenance are small, apparently, intrinsically disordered proteins, and several more are tiny membrane proteins. For these, *de novo* origin from non-coding sequence during virus evolution seems a likely option (53, 54). Pervasive transcriptional intitiation within poxvirus genomes could facilitate the generation of new proteins (55). The majority of the proteins in this category, however, appeared to be globular and yielded high quality AlphaFold2 models. Thus, it appears most likely that the core folds of these proteins have no counterparts among cellular proteins (or at least, these are extremely rare folds). Conceivably, these unique folds evolved by exaptation of host proteins that was accompanied by major rewiring of structural elements, resulting in unique topologies. A notable case in point in the family of poxvirus proteins containing the chemokine-binding PIE domain (47). In general shape, the PIE domain resembes other all-beta domains, such as the immunoglobulin fold, but the topology of the beta sheets is unique, so that structure comparisons detect no significant similarity. This apparent extensive protein fold remodeling suggests that evolution of viruses with large genomes is even more innovative than previously suspected. It has been observed that sequence diversity in a single family of large DNA viruses can surpass that of an entire domains of cellular life (56). The present observations complement these findings by demonstrating the commensurate structural diversity of virus proteomes.

## Methods

### Structure prediction of selected OPG representatives using AlphaFold2

For each OPG, a single member was chosen for structural model prediction with Alphafold2 2.2.0. Multiple sequence alignments (MSAs) were generated for each with the Alphafold2 pre-processing pipeline using default parameters and databases as of 2022-04-22. Exceptions were OPGs 174 and 189 which required additional hhblits parameters for the query against the BFD + Uniclust30 databases (OPG174: `-prepre_smax_thresh 50 -pre_evalue_thresh 100 -maxres 80000` ; OPG189: `-maxres 60000`). MSAs were used for structure modeling with the AlphaFold2 monomer model using a template date cutoff of 2022-01-01. Of the 5 models generated for each OPG by AlphaFold2 the model with the highest mean plddt was chosen for further analysis and structural similarity searches.

All AlphaFold2 models were assessed by their average and local plddt score. Of the 214 OPG models, 186 showed a reliable overall average plddt score of 70 or higher (Supplementary table S1). Additional 8 proteins contained one or more predicted globular domains with a local plddt score higher than 70. A single OPG (opg172) showed a globular fold but with a local plddt score below 70. The remaining 19 OPGs are short proteins for which single alpha helices can have high plddt scores but no globular fold was detectable. All models were kept for downstream analysis although results obtained from low quality models were further examined manually. In addition, proteins were classified into small (<=100 amino acids (aa)), intermediate (100-200 aa) and large (>200 aa). Ordered, globular stretches were identified as those with a plddt score of 70 or higher for 6 or more consecutive amino acids. All ordered stretches of a single protein were considered to classify the overall fold as either (partially) structured/globular (either at least 50% of the sequence, for small and intermediate proteins, or at least 100 amino acids, for large proteins, being part of ordered stretches) or disordered (failing the above criteria).

·

### Comparison of the OPG structural models to databases of protein structures

All high quality AlphaFold2 models of OPGs were compared to the pdb andAlphaFold2 (v2) databases using local installations of FoldSeek (2-8bd520) and Dali (5.1).

For foldseek prebuilt databases of PDB, AlphaFold/Proteome, and Alphafold/SwissProt were obtained with ‘ `foldseek databases`’ on 2022-06-04. Each structural model was used to query each of the three databases with ‘`foldseek search -s 9.5 --max-seqs 2000 -a`’ followed by conversion to html and tabular output files.

For Dali, a prebuilt database for AlphaFoldDB v2 was obtained from http://ekhidna2.biocenter.helsinki.fi/dali/AF-Digest.tar.gz. Before using this database, a small number of empty data files and structures with more than 200 structural elements had to be removed to accommodate limitations of Dali (989438 structures). A PDB database was built by importing a local mirror of PDB (507304 structures of subunits). For both databases, a 70% clustering was generated with ‘`cd-hit -c 0.7`’ to be used as a representative set in hierarchical searches with ‘`dali.pl --hierarchical –oneway –repset`’.

In order to compare all representative OPG structures with each other, a local Dali (12) all-vs-all run was performed (default settings, multimode mode with 50 nodes). Corresponing Dali Z-scores were visualized in an ordered matrix.

### Protein structure visualization

Protein structures were visualized with Chimera X (57). Superposition of proteins was either realized by Chimera X internal matchmaker (command ‘match #2 to #1) if protein sequences were similar enough. Otherwise, Dali translation-rotation matrices were used from the structural alignment runs. (Chimera X command ‘view matrix mod #2,’ followed by the 12 positions, comma separated, in order to match protein #2 to #1). See also: https://www.rbvi.ucsf.edu/pipermail/chimerax-users/2022-May/003656.html).

### Structural alignment

Structural alignments were generated using Dali web server (http://ekhidna2.biocenter.helsinki.fi/dali/) (12) by running the representative OPG pairwise against 7 diverse OPG members and 3 selected Dali hits, either from the pdb database (58) or from AlphaFold2 database (4, 59).

## Additional material

Additional files can be found at () and include:

AlphaFold2 models for all OPGs.

Structural alignments of the OPGs mentioned above and their respective homologues.

Dali all-vs-all Z-score matrix of representative OPGs.

## Author contributions

E.V.K., T.G.S. and B.M. initiated the project; E.V.K. designed research; W.R. and G.F. designed and ran the computational pipelines for protein structure modeling and comparison; P. M., T.G.S., E.V.K. and B.M. analyzed the results; P.M. performed structure superposition and alignment; E.V.K., P.M. and W.R. wrote the manuscript that was edited and approved by all authors.

## Acknowledgements

P.M. and E.V.K. are supported by the Intramural Research Program of the National Institutes of Health (National Library of Medicine). W.R. is supported by the Center for Information Technology, NIH; T.G.S. and B.M. were supported by the Division of Intramural Research, NIAID, NIH. This work utilized the computational resources of the NIH HPC Biowulf cluster. (http://hpc.nih.gov)

## Figure Legends

**Figure S1 (to Fig1):**
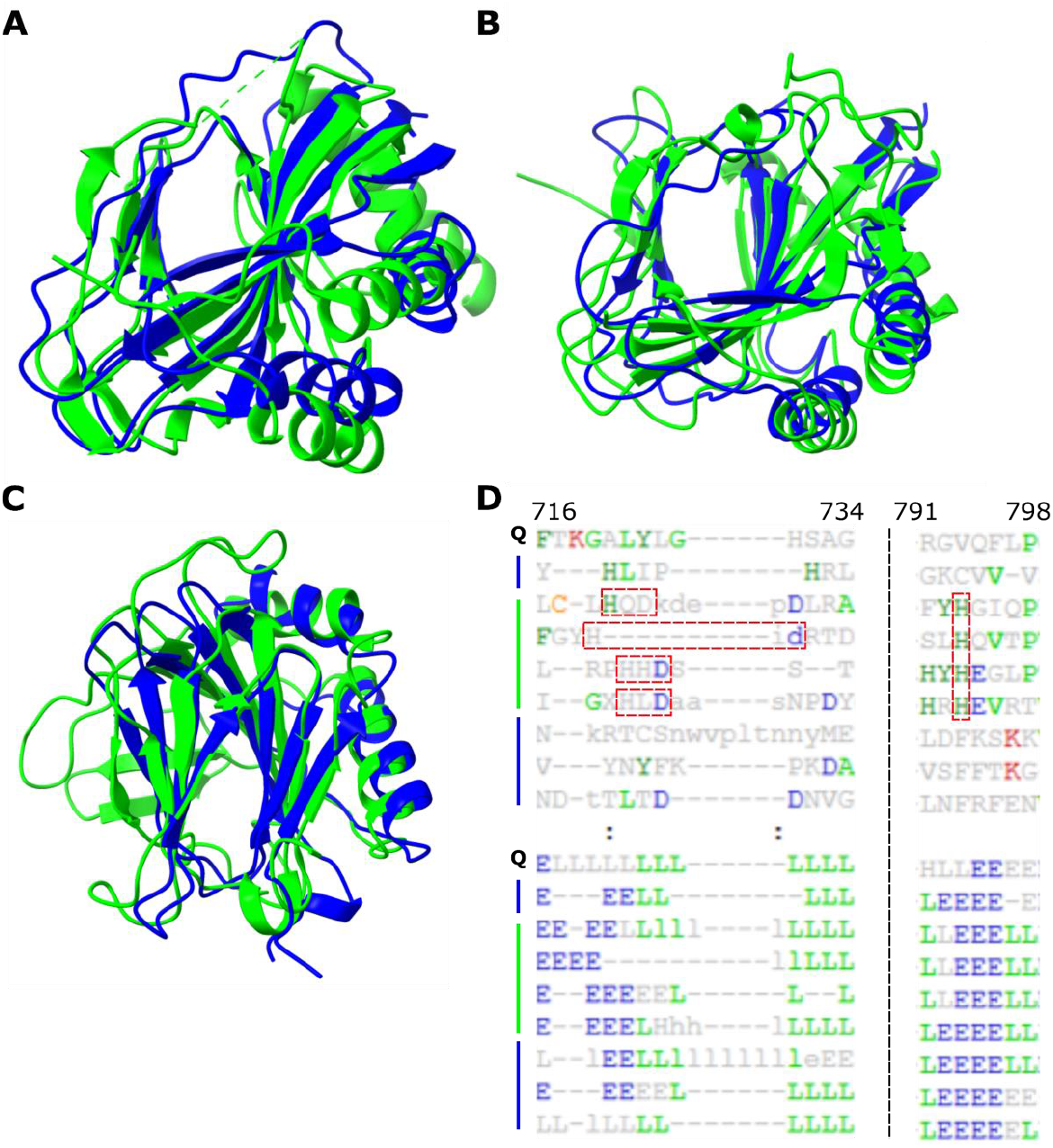
OPG20 (C10L), OPG31 (C4L) and OPG165 (A37R) show homology to hydroxylases. **A-C)** Superimposition OPG (blue) with respective hydroxylase domain of cellular protein (green). **A)** OPG20 (aa 1-160) and PKHD-type hydroxylase of Shewanella baltica (3dkq { DOI: 10.2210/pdb3dkq/pdb }, aa -5 – 200). **B)** OPG31 (aa 1-147) and Oxidoreductase, 2OG-Fe(II) Oxigenase family of Burkholderia pseudomallei (6n1f { DOI: 10.2210/pdb6n1f/pdb }, aa 3-216). **C)** OPG165 (aa 1-139) and dioxygenase from E. coli (3o1r (77), aa 13-214). **D)** Structural alignment of OPGs with hydroxylases as best hit. Green: OPGs from this figure and Fig.2 A and C with OPG55 (F11L) as query for pairwise structural alignment. From top to bottom: OPG55, OPG165, OPG20, OPG181 (A51R) and OPG31. Blue: hydroxylases, from top to bottom: dioxygenase from E.coli (3o1r (77)), PKHD-type hydroxylase of *Shewanella baltica* (3dkq { DOI: 10.2210/pdb3dkq/pdb }), human Lysyl Hydroxylase LH3 (6tex { DOI: 10.2210/pdb6tex/pdb }) and an oxidoreductase from *Burkholderia pseudomallei* (6n1f { DOI: 10.2210/pdb6n1f/pdb }).

**Figure S2:**
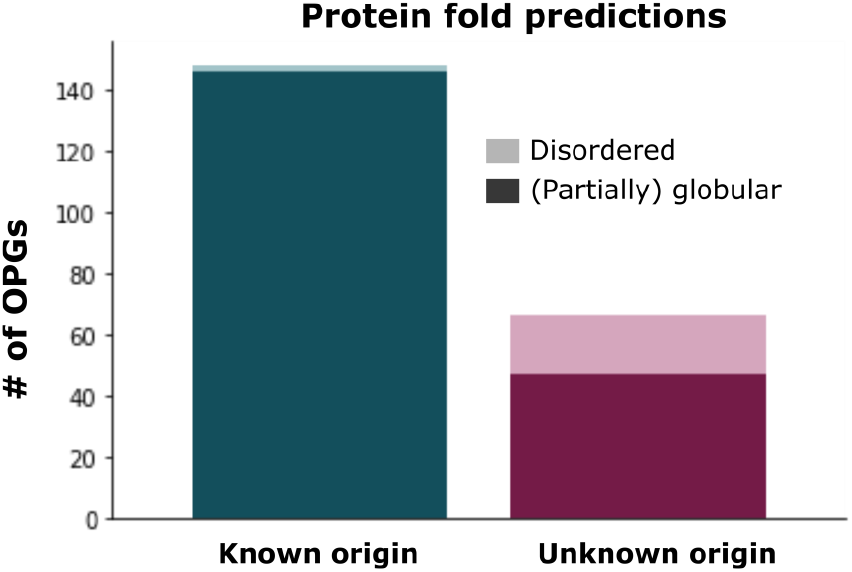
Predicted global folds of OPGs of known and unknown provenance. OPGs were classified as having a global disordered or (partially) globular fold based on their plddt score (see Methods for details). Average plddt for OPGs with known provenance is 86.1 (SD 8.9), for OPGs with unknown provenance 76.5 (SD 11.5) and for OPGs of unknown provenance with a globular fold 82.6 (SD 6.4). Excluding small OPGs with a generic fold from this fraction, the average plddt is 83.3 (SD 6.4).

**Figure S3:**
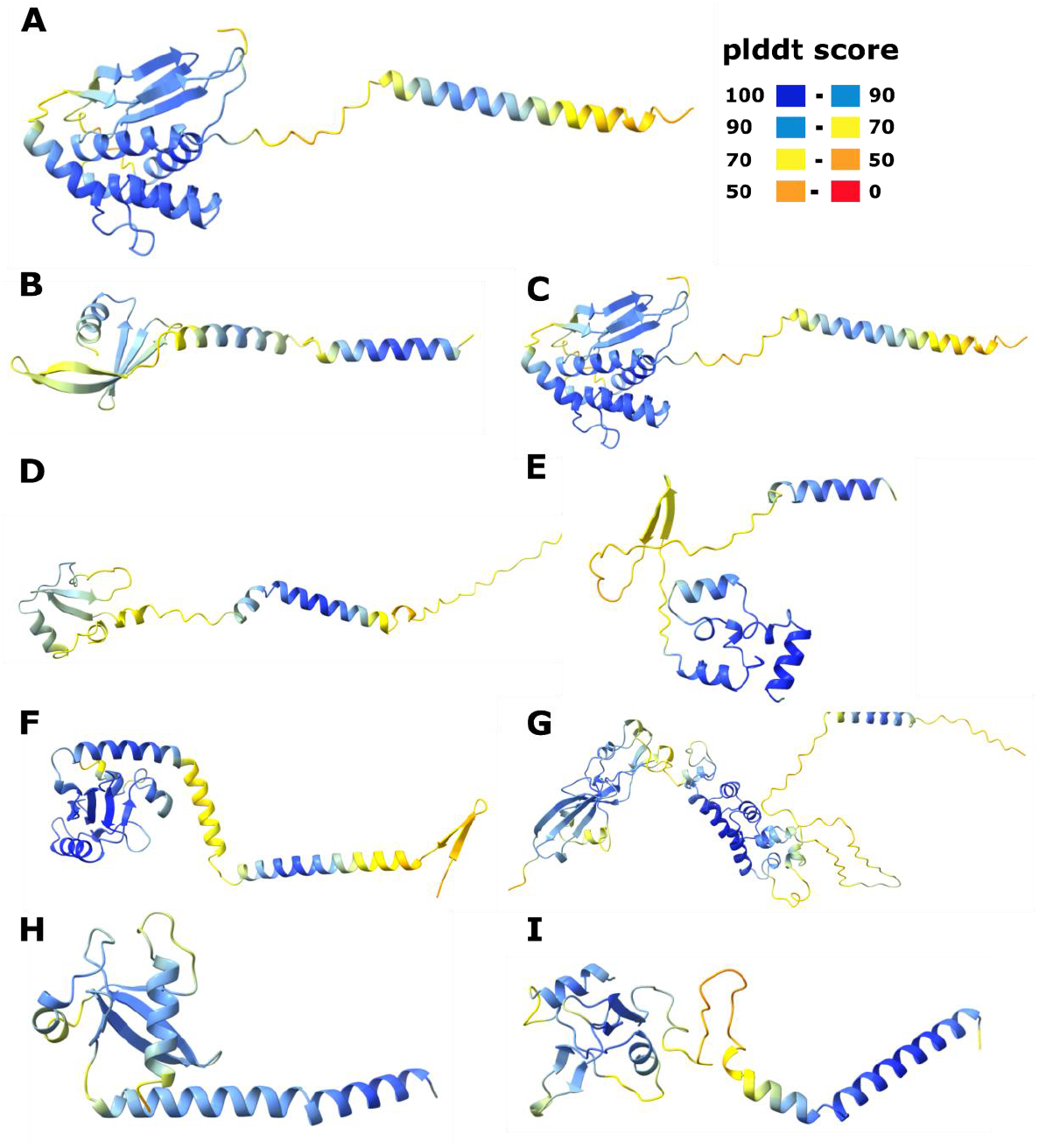
Predicted novel folds for OPGs including subunits of the viral entry-fusion complex. Predicted globular structures with no homologs detected outside of poxviruses are shown. A). **A)** OPG53 (F9L), **B)** OPG86 (G3L) **C)** OPG94 (G9R), **D)** OPG99 (L5R), **E)** OPG104 (J5L), **F)** OPG107 (H2R), **G)** OPG143 (A16L), **H)** OPG 147 (A21L) and **I)** OPG155 (A28L). The coloring is according to the AlphaFold2 plddt-score.

**Figure S4 (to Fig6):**
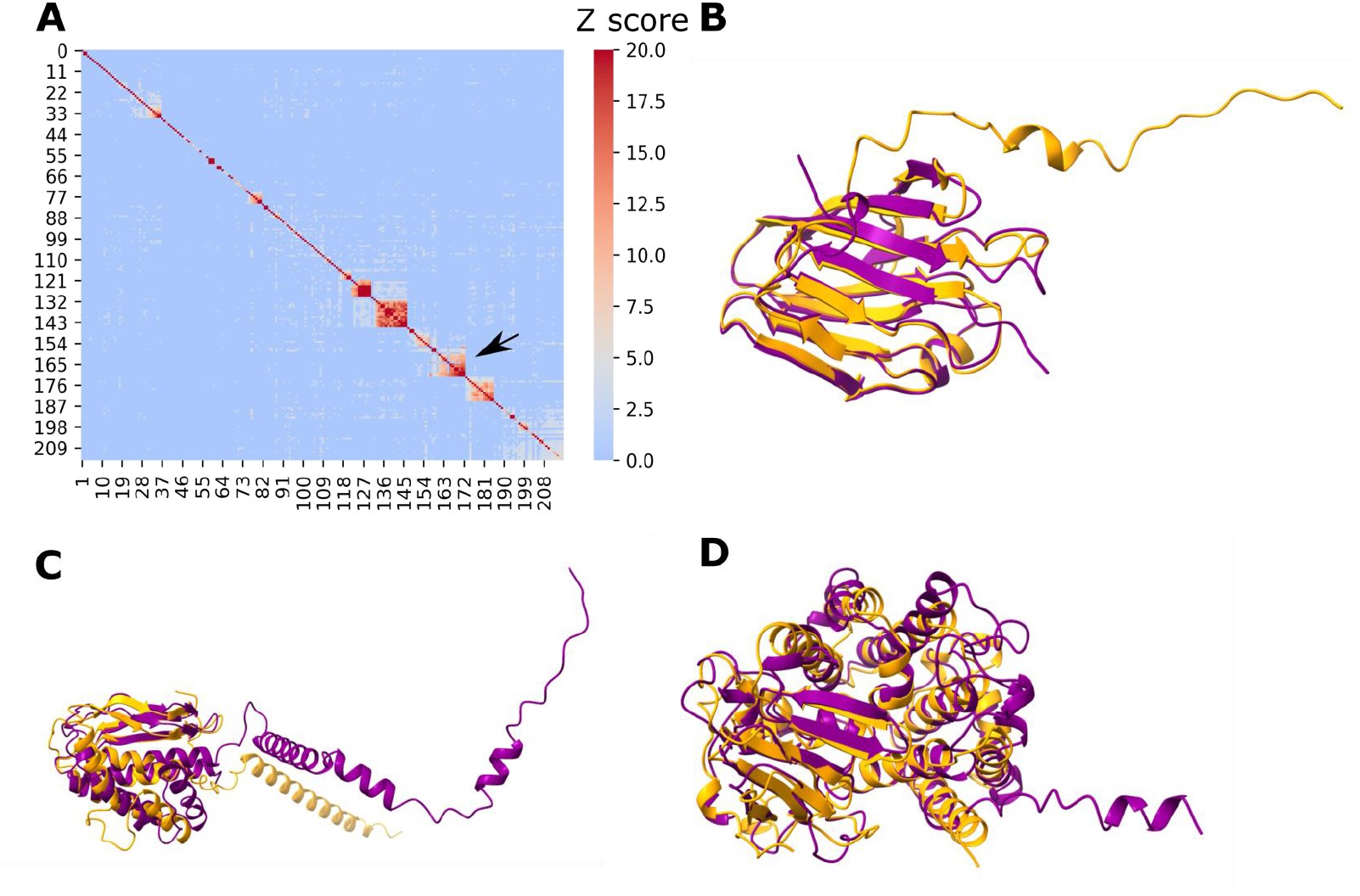
Recurring unique poxvirus protein structures across OPGs. **A)** Dali Z-score matrix for an all-vs-all run of all OPGs. OPGs were sorted based on Z-score, hence numbers at the axes are arbitrary. Arrow indicates the cluster of OPGs with PIE domains. For visualization, Z-score was caped at 20. Individual OPG pairs can have a Z-score beyond 20. **B-D)** Representative globular structures with no homologs identified outside of poxviruses but recurring among the OPGs. A) OPG27 (C7L) (purple) and OPG18 (orange), B) OPG95 (L1R) (purple) and OPG53 (F9L) (orange) and C) OPG153 (A26L) (purple, aa 1-359 (of 518) and OPG152 (A25L) (orange, aa 1-332 (of 1279))

### Abbreviations

VAVC: Vacinia virus
MPXV: Monkeypox virus
CMLV: Camelpox virus
VARV: Variola virus
SwPV: Swinepox virus
SORPV: Sea otterpox virus
LSDV: Lumpy skin disease virus
YLDV: Yaba-like disease virus
MyxV: Myxoma virus
MCV: Molluscum contagiosum virus
CRV: Nile Crocodilepox virus
CNPV: Canarypox virus
SGPV: Salmon gill poxvirus
SFV: Rabbit (shope) fibroma virus
af2-db: AlphaFold2 database
ECTV: Ectromelia virus
RCNV: Raccoonpox virus
YKV: Yoka poxvirus

